# Predictive vascular growth and remodeling in pulmonary hypertension: simulating intervention effects from captured evolution

**DOI:** 10.64898/2026.08.07.743318

**Authors:** Faeze Jahani, Brett Cardenas, Edward Manning, Jason M. Szafron

**Affiliations:** Department of Biomedical Engineering, Carnegie Mellon University, Pittsburgh, PA, US; Stanford Cardiovascular Institute, Stanford University School of Medicine, Stanford, CA, US; Section of Pulmonary, Critical Care and Sleep Medicine, Yale School of Medicine, Yale University, New Haven, CT, US

**Keywords:** Pulmonary hypertension, growth and remodeling, monocrotaline, therapeutic response, multiscale modeling

## Abstract

Pulmonary hypertension (PH) is characterized by progressive structural and mechanical remodeling of the pulmonary vasculature, yet few computational frameworks directly link disease mechanisms to longitudinal progression and therapeutic response. In this study, we utilized a multiscale pulmonary arterial growth and remodeling (G&R) framework to capture evolving functional metrics from rat models of PH. This framework couples morphometric tree hemodynamics, constrained mixture theory-based wall mechanics, and maladaptive cellular remodeling. Disease progression was driven by three mechanistically interpretable parameters governing excess smooth muscle production, remodeling activation, and passive stiffening. These parameters were calibrated to longitudinal monocrotaline (MCT) measurements of pressure, wall thickness, and stiffness from prior work using a multiobjective optimization. To show the predictive value of this model, we simulated therapeutic intervention within the same disease-specific framework by using functional cell-level responses to therapy to inform changes in parameter values. Calibration to the study-specific MCT dataset reproduced the temporal increases in pressure, wall thickness, and stiffness, demonstrating that the model could capture multiple features of vascular remodeling simultaneously, with *R*^2^ values of 0.81, 0.83, and 0.95, respectively,. Simulated treatment reduced pressure, wall thickness, and stiffness. Predicted pressure and wall-thickness responses agreed closely with the corresponding experimental treatment effects, whereas stiffness recovery was overpredicted, suggesting that additional mechanisms may contribute to persistent vascular stiffening after intervention. The framework also captured the overall progression of pulmonary pressure increases across both aggregated MCT and Sugen–hypoxia datasets, suggesting utility across studies and animal models. This work outlines a physics-based, multiscale framework that simulated quantities of direct clinical interest in a mechanistically interpretable platform for linking pulmonary vascular remodeling and treatment response. It supports comparisons across experimental phenotypes and interventions while identifying where constitutive refinements are needed to improve predictive capability across phenotypes.

## 1 Introduction

Pulmonary hypertension (PH) is a progressive cardiopulmonary disorder characterized by elevated pulmonary vascular resistance, increasing pulmonary arterial pressures, and eventual right ventricular dysfunction and failure [1]. Although PH is clinically defined by abnormal pulmonary hemodynamics, the disease is fundamentally driven by structural and mechanical remodeling of the pulmonary vasculature. Across PH subtypes, progressive luminal narrowing and structural stiffening alter vascular hemodynamics and increase right ventricular afterload over time [1–3]. A major clinical challenge is that disease progression and treatment response are highly heterogeneous, even within the same diagnostic group. Current clinical and preclinical tools provide valuable, repeated measurements of structure and function [4, 5], but these measures are not generally linked temporally with computational tools to characterize and predict outcomes. Thus, a major goal of this work is to demonstrate the utility of physics-based modeling tools to link data across measurement times and simulate quantities of clinical interest.

The pulmonary arterial wall provides a direct mechanobiological link between cell-scale remodeling and organ-level dysfunction. Compared with healthy vessels, remodeled pulmonary arteries in PH exhibit medial thickening, smooth muscle cell (SMC) proliferation, altered extracellular matrix (ECM) composition, and increased passive stiffness [1, 2, 6]. These changes increase vascular resistance and reduce compliance, thereby elevating right ventricular afterload and contributing to right ventricular remodeling. At the same time, changes in vascular geometry and wall composition alter the local mechanobiological cues experienced by endothelial cells, smooth muscle cells, and fibroblasts, which in turn modify subsequent growth and remodeling responses [2, 7, 8]. Therefore, pulmonary vascular disease emerges from a multiscale feedback process in which local cellular and matrix remodeling accumulates into network-level hemodynamic dysfunction. Indeed, altered mechanobiological cues can drive phenotypic changes that initiate inflammatory processes [9, 10]. As a result, PH progression is sustained not only by structural remodeling itself, but also by the self-amplifying interaction between altered mechanics and inflammation [11, 12].

Two mechanobiological stimuli are particularly important in pulmonary vascular remodeling: wall shear stress (WSS) and intramural stress (IMS). WSS arises from the frictional force exerted by flowing blood on the luminal surface and primarily regulates endothelial cell (EC) behavior. IMS results from pressure-induced deformation of the vessel wall and acts on SMCs and ECM constituents. Both stimuli are strongly dependent on vascular geometry and local hemodynamics [13, 14]. Because WSS exhibits a nonlinear dependence on vessel radius, even modest luminal narrowing can produce substantial changes in EC phenotype. In PH, elevated transmural pressure increases IMS, promoting SMC activation, changes in vascular tone, and ECM remodeling. It is well established that hemodynamic loading and mural composition varies across vessels of different sizes in the pulmonary vasculature [7]. Consequently, spatially heterogeneous changes across the vasculature can generate region-specific patterns of altered loading and remodeling in PH development [15, 16]. These observations motivate the use of a multiscale framework that couples hemodynamics, wall mechanics, and time-dependent inflammation across vessels of different sizes to best simulate PH evolution.

Rodent models provide an important platform for characterizing this progression in depth over time. Among the most widely used experimental models are monocrotaline (MCT)-induced and Sugen-hypoxia (SuHx) PH in rats, which span degrees of severity, mechanisms, and reversibility [17]. MCT induces a Group 1-like phenotype through endothelial injury and inflammation and is relatively reversible in rodents, particularly during early and intermediate stages. In contrast, SuHx combines vascular endothelial growth factor inhibition with hypoxia and produces a more severe, partially irreversible phenotype with distal vascular obliteration and advanced remodeling. Despite distinct triggers, both models converge on pulmonary arterial remodeling as a central disease driver, making them valuable for development of a unified vascular growth and remodeling framework. Longitudinal data from these models have also highlighted that vascular stiffening is an early and potentially pivotal regulator of pulmonary remodeling and pressure progression [4, 5, 18]. The ready availability of data from these models makes them an attractive target for use in developing novel biomechanical modeling tools to understand PH progression.

To capture evolving biomechanical data in animal models of vascular disease, constrained mixture theory (CMT)-based growth and remodeling (G&R) tools have been developed. In systemic hypertension, AngII-induced remodeling has been simulated with a single-vessel framework driven by prescribed pressure and immune cell activity [9, 19]. We previously extended the use of CMT G&R to capture the qualitative evolution of the pulmonary arterial tree in response to both adaptive and maladaptive perturbations to load or inflammatory stimuli [6]. Yet, we focused on qualitative remodeling behavior rather than direct calibration to longitudinal experimental data and relied on single parameter perturbations in a relatively large parameter space. Thus, formal numerical optimization of G&R parameters to time course data is necessary to demonstrate utility of these frameworks for real clinical simulations.

This study focuses on the integration of longitudinal data from animal models of PH into our previously developed G&R framework. To represent PH-related maladaptation, we focus on a subset of modeling parameters informed by the underlying functional mechanisms of inflammatory remodeling. We first demonstrate the sensitivity of key outputs from our framework to perturbations in these parameters, which motivates their use as variables in subsequent optimizations. We then identified parameter values that allow for matching to both aggregate time course data from the literature on single metrics and time course data on multiple metrics from a single prior study. After model calibration to the disease evolution, therapy is introduced through modification of inflammatory remodeling parameters according to changes observed experimentally in cell culture experiments. This allows predicted evolution of clinical metrics with intervention and comparison to actual outcomes, along with parametric studies of functional effects that could guide hypothesis development and further treatment approaches. Overall, this shows our ability to develop, calibrate, and evaluate a predictive pulmonary arterial G&R model to represent disease progression and intervention response informed by multiple experimental data sets.

## 2 Methods

### 2.1 Multiscale G&R framework for PA tree simulations

Our previously developed multiscale pulmonary arterial G&R framework was used to simulate vascular remodeling during PH. A detailed description of the model structure and development were previously published [6], and key aspects are summarized here. This model couples a Strahler-diameter defined morphometric tree for hemodynamic modeling with a CMT-based description of the vessel wall throughout the vascular network [6]. At each remodeling time step, the framework (i) generates or updates the pulmonary arterial tree, (ii) solves network hemodynamics, (iii) computes loaded vessel geometry and local mechanical quantities across vessels of different caliber, (iv) evaluates mechanobiological and inflammatory stimuli, (v) updates production, removal, and material behavior of each structurally-significant constituent, (vi) performs fixed-point iterations to reach convergence of G&R and hemodynamic changes at each time step and (vii) advances the system forward to the next discrete time point for update of the entire system starting again from (i). This structure enforces mechanical consistency at each time point while capturing disease progression over weeks.

The pulmonary vasculature morphometric arterial tree was derived from measured pulmonary geometries [20]. For each vessel order, representative values of radius, wall thickness, and length were assigned, allowing remodeling to be simulated throughout the network in a computationally efficient way. For each simulation, blood flow and pressure were solved throughout the network using conservation of mass at bifurcations and Poiseuille-type segmental resistance relationships [21]. These calculations provided local hemodynamic quantities for each vessel order, including pressure, flow, WSS, and IMS [6]. Hemodynamic loads were then passed to the G&R model, where deviations from homeostatic stresses and the current inflammatory burden generated remodeling stimuli. As vessel geometry and material properties evolved over time, the hemodynamics were recomputed, producing a closed feedback loop between local wall evolution and network-level function.

The vessel wall was represented as a constrained mixture of structurally important constituents, including elastin and ground matrix, collagen, and SMCs [22]. Baseline mechanical behavior of the pulmonary arterial wall was established from biaxial mechanical testing of healthy large PAs and an optimization to distribute heterogeneous behavior across the vasculature at the initial time point to maintain the specified loaded geometry from the morphometric data [4, 6]. After establishment of the initial state, each constituent contributed to the overall wall response over time through its own material behavior, production rate, degradation kinetics, and deposition history. Although the framework accounts for multiple wall constituents, the present study focused primarily on smooth muscle cell remodeling as a key mechanism underlying the disease-specific remodeling responses considered here.

### 2.2 Inflammatory remodeling relations driving PH development

Disease-induced pulmonary vascular remodeling was represented through maladaptive inflammatory mechanisms that increase smooth muscle production and alter passive smooth muscle material behavior [12, 23, 24]. The three key disease parameters calibrated in this study were the inflammatory gain on SMC proliferation *K_i_*, the time constant for inflammatory development *δ_i_*, and the passive SMC stiffening factor *γ_i_*. In this formulation, *K_i_* controls amplitude of the inflammatory response kinetics, *δ_i_* modulates the timeline for disease development, and *γ_i_* affects material behavior changes due to remodeling rather than mass changes. We describe the constitutive relations for SMCs below containing these parameters, while formulations for other constituents were described previously [6]. For the uniform-inflammation prescribed herein across vessels, the baseline maladaptive remodeling parameters were prescribed as

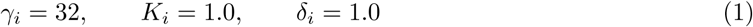

These values were used as the reference parameter set for perturbation in parametric studies prior to fitting time course data. Within the modeling framework, *K_i_* is defined within the equation describing the SMC production rate *m*(*s*) at remodeling time *s* as

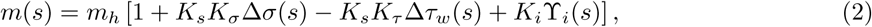

where *m_h_* is the homeostatic smooth muscle production rate, *K_s_* modulates the adaptive mechanobiological sensitivity, *K_σ_* and *K_τ_* weight the effects of intramural stress and wall shear stress, respectively, and *K_i_* indicates the specific application of inflammatory remodeling to SMCs. The time varying inflammatory stimulus function Υ*_i_*(*s*) was given by,

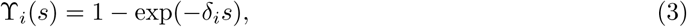

where *δ_i_* controls the rate of inflammatory onset. Functional forms for stored energy density of SMCs were also modified to allow for passive stiffening associated with intracellular reorganization and phenotypic change. The inflammatory increment in passive SMC stored energy was written as

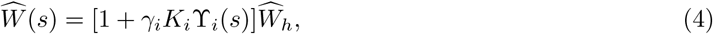

where *Ŵ_h_* is the homeostatic passive SMC strain-energy contribution and *γ_i_* governs the magnitude of passive stiffening or phenotypic change associated with maladaptive remodeling [6]. All other equations and parameter values remain fixed to those used previously, as we sought to limit our focus to parameters that are easily interpretable. To facilitate code sharing and use of external libraries, the codebase for coupling G&R simulations to hemodynamics was converted to Python for this and verified against prior results (Supplemental Figure 9). Our modeling framework is available in an online GitHub repository. To evaluate how these maladaptive remodeling parameters influence disease progression, we performed parametric studies with changes in individual parameter values while all others were held constant. The parameters *K_i_*, *δ_i_*, and *γ_i_* were varied over prescribed ranges from the baseline values used previously, and the resulting changes in pressure, wall thickness, and stiffness were evaluated over time. For *K_i_* and *γ_i_*, incremental linear changes were used, as order of magnitude changes led to poor model convergence, such as unbounded decreasing diameters for high degrees of inflammation. For *δ_i_*, order of magnitude changes were required to affect alteration of the time course, as only the onset of inflammation shifted rather than the magnitude. Across metrics, parametric studies were used to show that these functional pathways strongly affect pulmonary vascular remodeling and to interpret how each parameter contributed to specific changes in output metrics.

### 2.3 Parameter optimization and model building

A major goal of this work is to parameterize our underlying G&R model to capture real, time course data from studies on PH progression. Thus, we gathered data from the literature on two commonly used animal models of PH, MCT and SuHx-induced, which represent distinct disease etiologies. The most common data with a direct correlate to our model outputs was right ventricular systolic pressure (RVSP) as a direct measure of cardiac afterload and indirect measure of vascular remodeling. Aggregated data for pressures were extracted and normalized to the average week 0 pressure from multiple studies for both the MCT (13 studies from 0 to 4 weeks, Supplemental Table 2) and SuHx animal models (6 studies from 0 to 8 weeks, Supplemental Table 3). Amongst these studies, Liu et al., 2016 included additional data on distal vascular thickness and stiffness, with weekly values from 0 to 4 weeks in MCT rats and 5 time points from 0 to 8 weeks in SuHx rats [18]. This offered additional targets for direct matching from our G&R framework, as thickness is calculated as part of the equilibrium solution for the loaded geometry of each vessel order and stiffness can be derived from the material behavior according to a small-on-large approach to linearization of the current configuration mechanical behavior [25]. Additionally, this work included comparable data with administration of prostacyclin analogs in both cell culture and animal models, which offered an opportunity to make predictive simulations dependent on treatment response. Table 1 summarizes the successive modeling performed with these data.

**Table 1:** Simulations performed across literature datasets.

| Dataset | Disease model | Time course | Primary measurements | Role in study |
| --- | --- | --- | --- | --- |
| Aggregated MCT | MCT-induced PH | 4 weeks | Mean pressure progression across studies | Demonstrate fitting of time course data |
| Aggregated SuHx | SuHx-induced PH | 8 weeks | Mean pressure progression across studies | Illustrate difficulties in parameter estimation |
| MCT (study-specific) | MCT-induced PH | 4 weeks | Pressure, wall thickness, passive stiffness | Fitting of inflammatory parameters to equal number metrics |
| SuHx (study-specific) | SuHx-induced PH | 8 weeks | Pressure, wall thickness, stiffness | Evaluation of fitting method in additional dataset |
| Therapy intervention data | MCT-induced PH + treprostinil treatment | MCT 0–2 weeks, treprostinil 2–4 weeks | Pressure, wall thickness, stiffness | Prediction of treatment response without optimization |

These measurements provided additional targets for direct comparison with the G&R framework, as wall thickness is obtained from the equilibrium solution for the loaded geometry of each vessel order, while stiffness is derived from the current-configuration material response using a small-on-large linearization approach [25]. In Liu et al. [18], MCT-induced PH was initiated in adult male Sprague–Dawley rats using a single subcutaneous dose of MCT (50 mg/kg). For the treatment study, treprostinil was initiated 2 weeks after MCT administration and delivered intravenously at 90 ng/kg/min for an additional 2 weeks using a subcutaneous minipump connected to a jugular-vein cannula. In the SuHx model, rats received a single subcutaneous dose of SU5416 (20 mg/kg), followed by 3 weeks of exposure to 10% oxygen and 5 weeks of normoxic recovery, for a total study duration of 8 weeks. These experimental protocols provided longitudinal disease and treatment-response data for model calibration, evaluation, and predictive simulations. Table 1 summarizes the modeling analyses performed using these data.

As shown in Fig. 1, model parameters were estimated by minimizing the mismatch between simulated and experimental time-course data. Optimization was performed using the Nelder–Mead method [26], which iteratively updated the candidate parameter set to reduce the total normalized mismatch. Depending on the simulation, the optimized parameter set included a combination of *K_i_*, *δ_i_*, and *γ_i_*, which govern the sensitivity, onset, and passive mechanical consequences of maladaptive remodeling. For each candidate parameter set, the coupled morphometric tree hemodynamics and G&R model was simulated forward in time and the predicted outputs were compared against available experimental measurements depending on the dataset utilized in a given simulation.

**Fig. 1:**
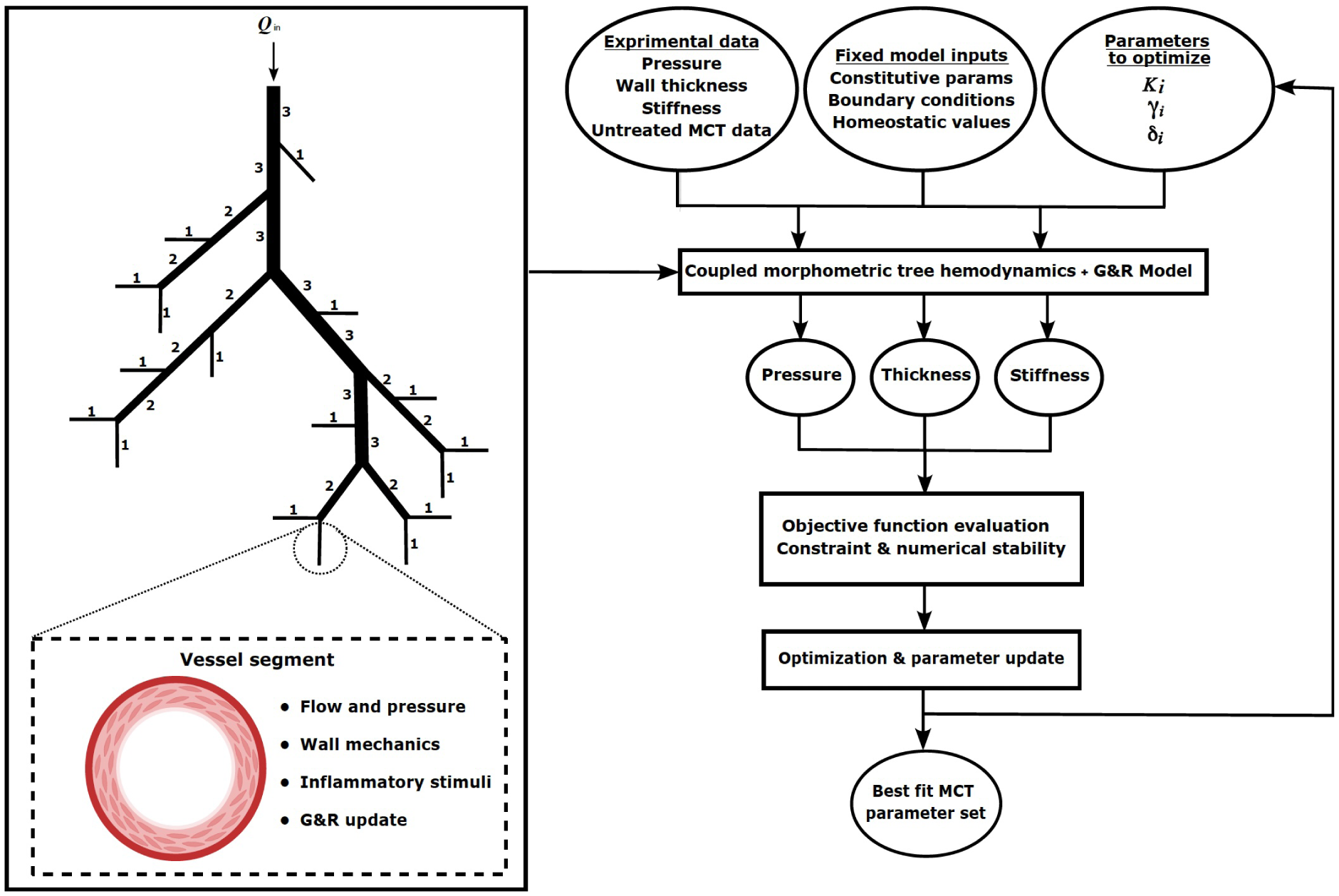
Parameter-optimization framework for the coupled morphometric tree hemodynamics and G&R model. **Left:** multiscale morphometric tree representation of the pulmonary vasculature, with prescribed inlet flow and outlet pressure used to solve network hemodynamics and pass mean pressure and flow to the G&R model for geometry and remodeling updates. **Right:** optimization workflow in which fixed model inputs, candidate maladaptive remodeling parameters, and experimental data are used to predict pressure, wall thickness, and stiffness; these outputs are compared with measurements through objective-function evaluation and feasibility constraints to identify the best fit disease parameter set.

The objective function was defined as the normalized, mean-squared error between the simulated and experimental time course. Through our initial parametric studies, physiologically realistic bounds were determined, with 0 *≤ K_i,_*_steady_ *≤* 16, 0 *≤ δ_i_ ≤* 1, and 0 *≤ γ_i_ ≤* 48, that allowed for convergence and demonstrable effects on the output metrics of interest. However, in optimization across multiple parameters, simulations could produce non-physiologic states or cause solver failure with unbounded narrowing or dilatation driven by excessive inflammation. In these cases, the lack of convergence was indicated as an infeasible objective evaluation. The optimization loop updated the candidate parameter set iteratively until the simulation data mismatch was minimized, which produced the calibrated parameter sets.

To reduce sensitivity to local minima, Latin Hypercube Sampling (LHS) [27] was used to generate diverse initial guesses within the acceptable parameter ranges for the deterministic Nelder-Mead algorithm. LHS acts across each parameter range in a structured way, so the initial candidate sets evenly cover the parameter space rather than clustering together. These sampled initial guesses were then used as starting points for repeated optimization runs. Convergence of multiple runs to the same or similar parameter values was interpreted as evidence of a stable optimum, whereas large spread among converged solutions indicated potential non-identifiability.

### 2.4 Therapeutic intervention simulations

After calibration of the MCT disease model, therapeutic intervention was introduced by modifying the maladaptive inflammatory remodeling parameters at the time of drug administration. Therapy was represented as a gradual, time-dependent reduction in maladaptive smooth muscle production and passive stiffening. These changes were incorporated into the vessel-level G&R equations and subsequently affected whole-tree geometry and hemodynamics through the coupled remodeling feedback. The intervention parameters were informed directly by experimental measurements rather than optimized, thereby allowing the predictive capability of the calibrated framework to be evaluated.

The simulations were based on the treprostinil experiment reported by Liu et al. [18]. Male Sprague– Dawley rats received MCT, and treprostinil treatment was initiated 2 weeks later at 90 ng/kg/min and continued for an additional 2 weeks. The same study also reported the effects of iloprost, a prostacyclin analogue similar to treprostinil, on pulmonary arterial smooth muscle cell proliferation and traction force in culture. These cell-level measurements were used to determine the treatment-induced reductions in maladaptive smooth muscle production and passive stiffening.

To constitutively model treatment response, we introduced

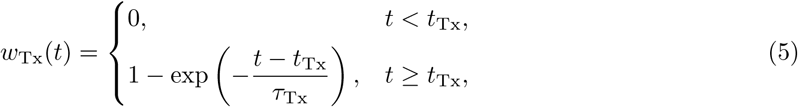

where *t*_Tx_ = 14 days is the treatment-initiation time and *τ*_Tx_ controls the time scale over which the treatment becomes effective. Thus, *w*_Tx_(*t*) describes the activation of the therapeutic effect.

The treatment-modified maladaptive production gain was defined as

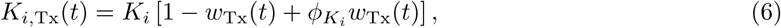

and the treatment-modified passive stiffening gain was defined as

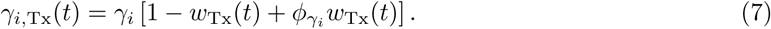

Here, *ϕ_K_i__* and *ϕ_γ_i__* represent the fractions of maladaptive smooth muscle production and passive stiffening, respectively, that remain under treatment, with smaller values corresponding to stronger treatment effects. Based on the reductions in pulmonary arterial smooth muscle cell proliferation and traction force measured following iloprost treatment in Liu et al. [18], these values were prescribed as *ϕ_K_i__* = 0.65 and *ϕ_γi_* = 0.23. A value of *τ*_Tx_ = 1 day was selected to represent a physiologically reasonable treatment onset.

The inflammatory production stimulus entering the vessel-level G&R equations was therefore written as

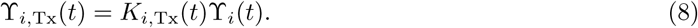

Passive smooth muscle stiffening was modified through modulation of the SMC stored energy

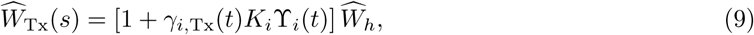

where *W_h_* is the corresponding homeostatic stored energy density. In this formulation, treatment modifies maladaptive smooth muscle production through *K_i,_*_Tx_(*t*) and passive material behavior through *γ_i,_*_Tx_(*t*). The baseline disease activation Υ*_i_*(*t*) remains unchanged, so the therapeutic effect is introduced only once through *w*_Tx_(*t*).

The resulting changes in vessel radius, wall thickness, and stiffness were passed back to the morphometric tree model, where whole-tree hemodynamics were recomputed. The therapy framework therefore used the best-fit untreated disease model as its baseline, applied experimentally informed reductions in maladaptive production and passive stiffening after treatment initiation, and predicted treated trajectories of pressure, wall thickness, and stiffness. The treatment-prediction workflow is summarized in Fig. 2, which illustrates how the calibrated disease parameters are combined with therapy-specific inputs to predict outcomes relevant to right ventricular systolic pressure, pulmonary compliance, and right ventricular function.

**Fig. 2:**
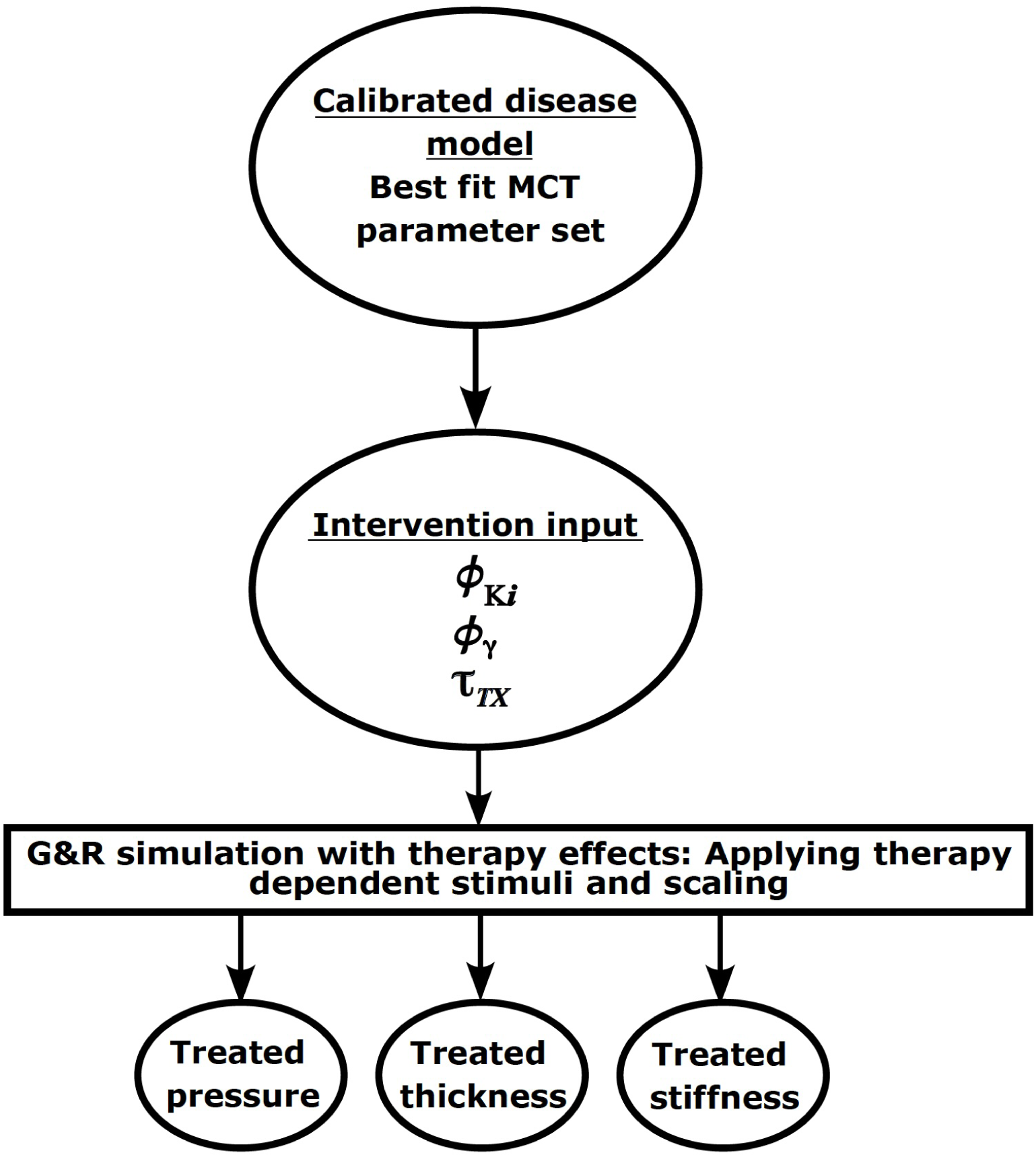
Therapy-prediction framework built on the calibrated disease model. The best-fit disease parameter set from optimization is combined with therapy inputs, including target fractions for maladaptive production and stiffening and the therapy activation timescale. The therapy-modified G&R simulation predicts treated pressure, wall thickness, and stiffness.

## 3 Results

### 3.1 Parametric study of inflammatory remodeling parameters

We first performed parametric studies to isolate the effects of the principal inflammatory remodeling parameters *K_i_*, *γ_i_*, and *δ_i_* on disease progression. Each parameter was varied independently while the remaining parameters were held fixed, and model outputs were reported as fold changes in pressure, wall thickness, and stiffness relative to baseline. As shown in Fig. 3, increasing *K_i_* increased all three outputs, with particularly strong effects on pressure and stiffness. Wall thickness also increased with *K_i_*, although its trajectories showed a more limited dynamic range and a tendency toward plateauing at later times. In contrast, increasing *γ_i_* most strongly amplified stiffness, produced a moderate increase in pressure, and had only a relatively small effect on wall thickness. This pattern is consistent with the role of *γ_i_* in controlling passive stiffening rather than growth itself. Notably, cross-over between simulations was identified in thickness evolution, where values of *γ_i_* led to lower final values, highlighting the potential non-linear and non-monotonic effects of these parameter changes on particular metrics. The parameter *δ_i_* primarily controlled remodeling onset and progression rate. Larger *δ_i_* values produced earlier and steeper increases in pressure, thickness, and stiffness, whereas smaller values delayed the response and reduced fold changes over the simulated time window. This timing effect was most evident in pressure and stiffness. Again, non-linear changes with perturbation were evident. End simulation values converged to similar levels for the most rapid infiltrations, suggesting that ultimate magnitude is minimally affected by varying *δ_i_* alone.

**Fig. 3:**
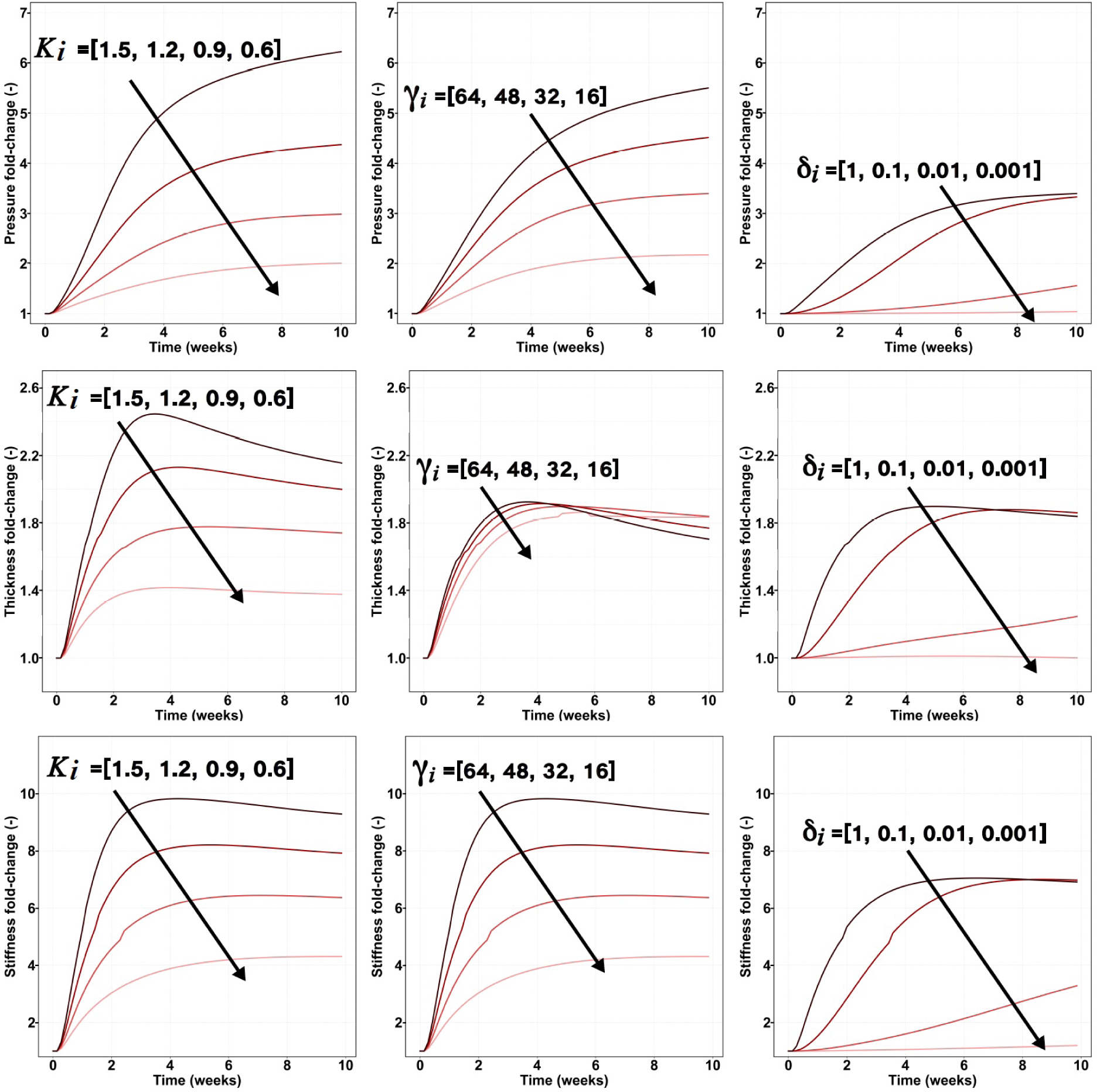
Parametric study of the principal inflammatory remodeling parameters. Fold-change trajectories of pressure (top), wall thickness (middle), and stiffness (bottom) are shown for independent variation of *K_i_* (left), *γ_i_* (center), and *δ_i_* (right). Increasing *K_i_* elevates all three outputs, increasing *γ_i_* primarily amplifies stiffness, and decreasing *δ_i_* delays and attenuates remodeling over the simulated time window.

### 3.2 Pressure progression across aggregated datasets

Next, we sought to validate the ability of our modeling framework to accurately capture the real time course of PH development from disease. We optimized our inflammatory parameters to capture the mean temporal progression of pulmonary pressure across aggregated MCT (Supplemental Table 2) and SuHx (Supplemental Table 3) datasets. To assess the importance of multi-parameter optimization, we first fitted the data using *K_i_* alone and then progressively included *δ_i_* and *γ_i_*, resulting in one-, two-, and three-parameter optimization schemes. The reduced-parameter fits are shown in Supplemental Fig. 10. The MCT pressure trajectory was captured well using *K_i_* alone, whereas the SuHx trajectory was poorly represented by the one-parameter model and improved as additional parameters were included, motivating the use of the full three-parameter optimization.

Building on the reduced-parameter analysis, we next evaluated the final model fits obtained by optimizing all three inflammatory remodeling parameters. As shown in Fig. 4, with all three inflammatory parameters the model captured the increasing pressure fold-change in both disease models while preserving their distinct progression patterns. For aggregated MCT data (Fig. 4A), the model captured the pressure trajectory over 4 weeks with *R*^2^ = 0.9782 and RMSE = 0.0606. For aggregated SuHx data (Fig. 4B), the model captured the pressure trajectory over 8 weeks with *R*^2^ = 0.9046 and RMSE = 0.2770. Relative to MCT, the SuHx datasets exhibited a greater magnitude of pressure elevation and higher inter-study variability. Together, these results support the robustness of the framework across heterogeneous datasets and its applicability across PH phenotypes.

**Fig. 4:**
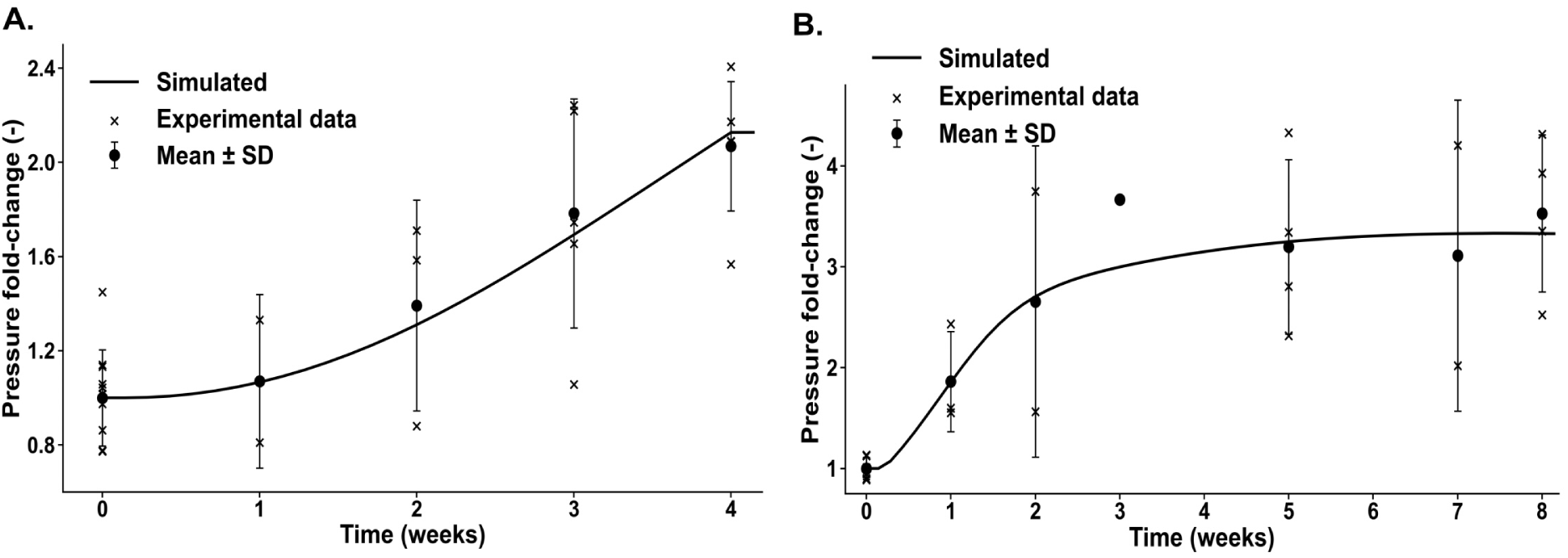
Pressure progression across aggregated pulmonary hypertension datasets. **(A)** Aggregated MCT data and corresponding simulated pressure fold-change over 4 weeks. **(B)** Aggregated SuHx data and corresponding simulated pressure fold-change over 8 weeks. Solid lines denote simulated trajectories, crosses denote individual experimental data points, and filled circles with error bars denote experimental mean *±* SD at each time point.

### 3.3 Study-specific multi-objective optimization

We next calibrated the model to the Liu et al. [18] MCT dataset by fitting pressure, wall thickness, and stiffness simultaneously over the 4-week study period. As shown in Fig. 5, the optimized model reproduced the progressive increase in all three outputs, while preserving the distinct temporal patterns of each metric. Pressure showed a nonlinear upward trajectory with increasing mismatch at later times, wall thickness increased more gradually and was captured with the smallest absolute error, and stiffness exhibited the strongest overall agreement across the full time course. Quantitatively, the optimized model achieved *R*^2^ = 0.8089 and RMSE = 0.3077 for pressure, *R*^2^ = 0.8309 and RMSE = 0.0691 for wall thickness, and *R*^2^ = 0.9476 and RMSE = 0.5983 for stiffness. Among the three outputs, stiffness was captured most accurately, whereas pressure and wall thickness showed slightly larger residual mismatch. Overall, these results indicate that the inflammation-driven G&R framework can simultaneously represent pressure elevation, wall thickening, and passive stiffening in MCT-induced PH. Optimization robustness was assessed using 15 initial parameter sets generated by Latin hypercube sampling [27]. Nine optimization runs converged to similar solutions, whereas six reached poorer local optima. The corresponding optimization trajectories and convergence behavior are shown in Supplemental Fig. 11. The corresponding best-fit parameter values, incorporated into Eqs. (2) and (4), were

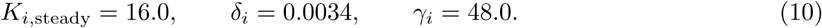

**Fig. 5:**
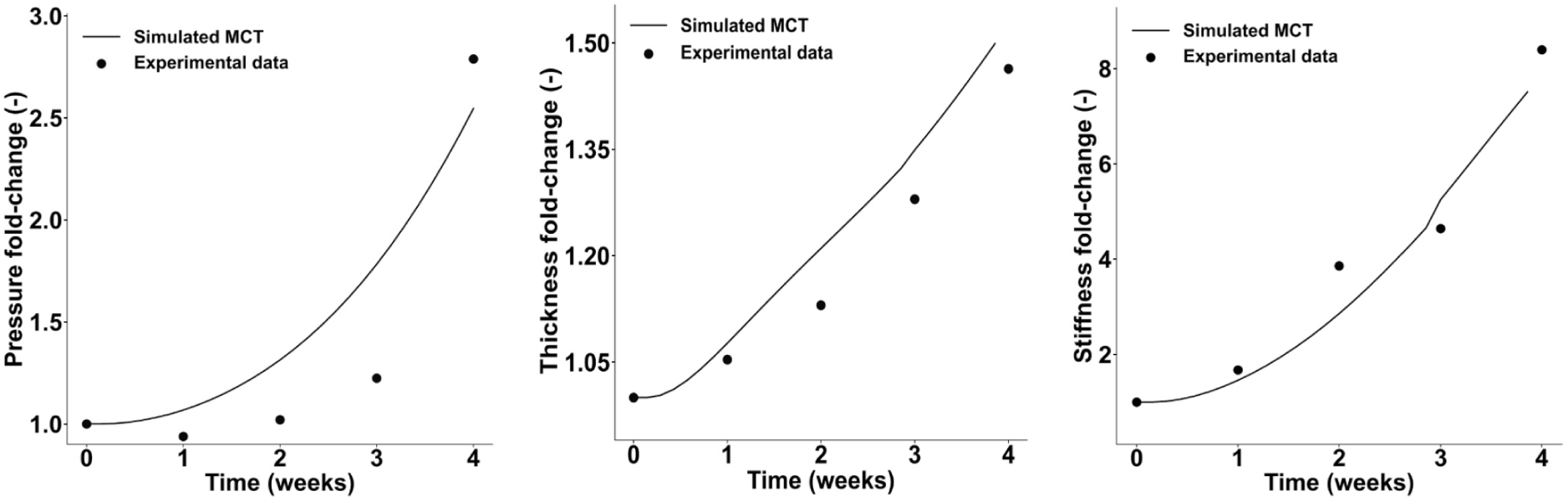
Study-specific multi-objective optimization for MCT dataset. Simulated and experimental fold changes are shown for pressure (left), wall thickness (center), and stiffness (right). The optimized model captures the progressive increase in all three metrics over time, with the strongest agreement observed for stiffness.

Notably, *K_i,_*_steady_ and *γ_i_* reached their prescribed upper bounds of 16 and 48, respectively, whereas *δ_i_* = 0.0034 was near the lower end of its allowable range of 0 to 1. Because smaller values of *δ_i_* correspond to a slower development of the inflammatory remodeling response, these results suggest that the experimental data favored strong maladaptive smooth muscle production and passive stiffening combined with relatively slow remodeling activation. Overall, these optimized values demonstrate that the framework can capture MCT-induced PH and reproduce the overall trajectory of disease progression.

Model performance was further assessed using a similar SuHx dataset reported by Liu et al. [18]. As shown in Fig. 6, the model poorly captured the trends in pressure, wall thickness, and stiffness with weaker agreement than for the corresponding MCT study-specific fit. The resulting RMSE values were 0.7470 for pressure (2.5X that from MCT fitting), 0.1757 for wall thickness (3X that from MCT fitting), and 0.5870 for stiffness (similar to that from MCT fitting). While the framework captured the general increases of these parameters in SuHx remodeling, the underlying remodeling formulation likely requires more detailed constitutive relations with more temporal parameters to better represent the mechanisms driving this more severe phenotype.

**Fig. 6:**
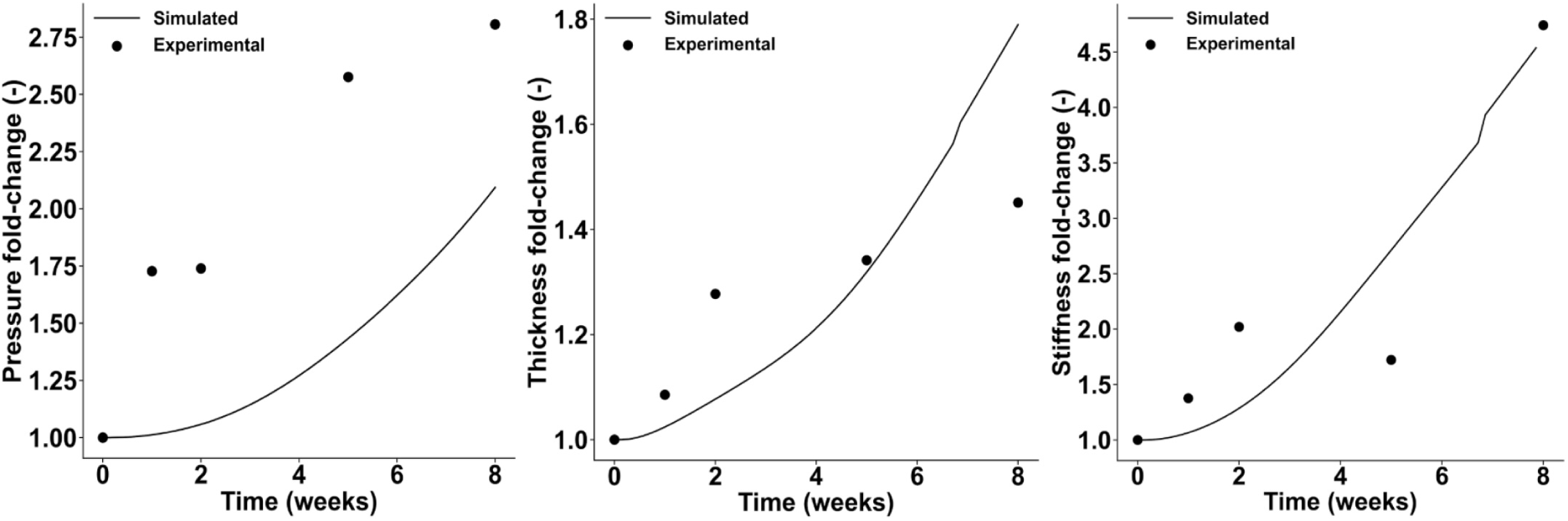
Study-specific evaluation in SuHx dataset. Simulated and experimental fold-changes are shown for pressure (left), wall thickness (center), and stiffness (right). The model captured the overall increasing trends in all three outputs, although with weaker agreement than in the corresponding MCT study-specific fit.

### 3.4 Prediction of treatment response in MCT-induced PH

After calibrating the MCT disease model with good matching to the observed progression, we sought to determine if our modeling framework could predict the effects of therapy from functional changes observed in cell culture. Therapeutic effects were introduced through Eqs. (6), (7), and (8), with therapy onset prescribed at *t*_therapy_ = 14 days, as performed in Liu et al. [18]. Consistent with the experimental observations of functional changes to SMC proliferation and traction development, our formulation represented a time dependent reduction in maladaptive smooth muscle production and passive stiffening after disease initiation. As shown in Fig. 7, the calibrated model with inclusion of therapy predicted partial reversal of remodeling, with reductions in pressure, wall thickness, and stiffness relative to untreated MCT. At week 4, the untreated simulation showed changes in pressure, wall thickness, and stiffness of 2.55, 1.53, and 7.51 fold, respectively, whereas the corresponding treated values were 1.48, 1.19, and 3.05. These changes correspond to simulated reductions of 41.92% in pressure, 22.17% in wall thickness, and 59.41% in stiffness.

**Fig. 7:**
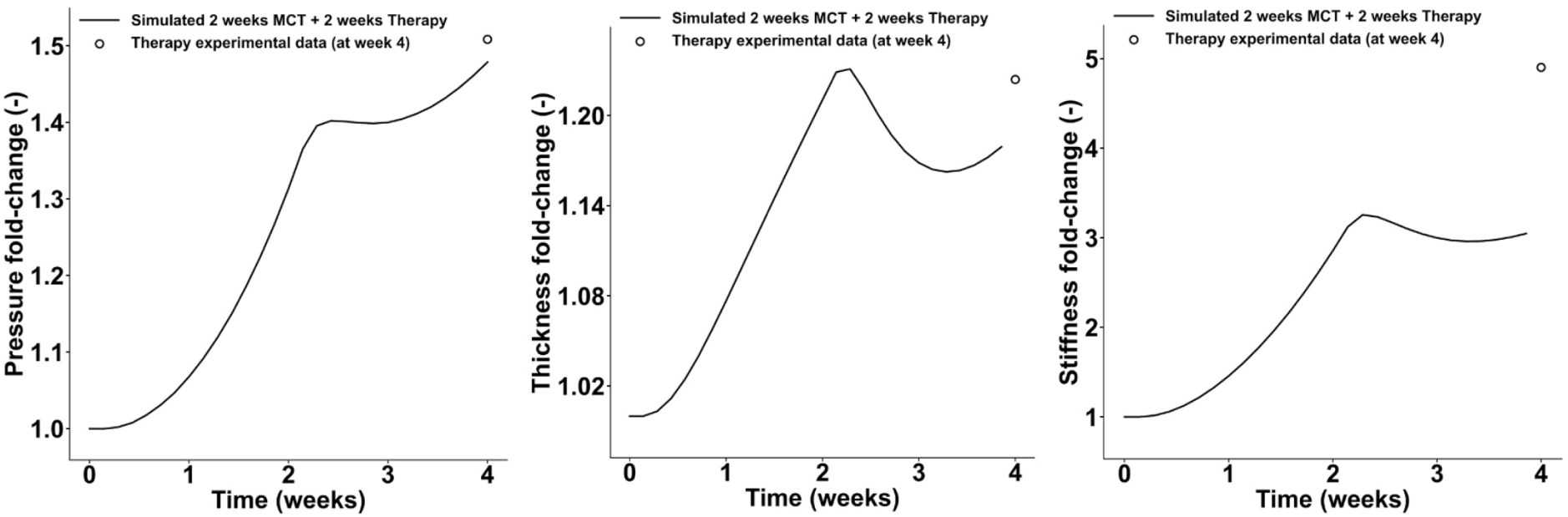
Prediction of treatment response in MCT-induced PH. Simulated treated trajectories are shown for pressure (left), wall thickness (center), and stiffness (right), with week-4 experimental therapy points mapped to the same fold-change scale for comparison. Therapy reduced all three outputs relative to untreated MCT, with close agreement for pressure and wall thickness and larger residual mismatch for stiffness.

To compare treatment predictions with the reported intervention data from rat models in the same study, the experimental 4-week therapy values were mapped to the same fold-change scale as the simulated MCT trajectories. The resulting experimental comparison points were 1.51 for pressure, 1.22 for wall thickness, and 4.90 for stiffness, corresponding to reductions of 40.75%, 19.80%, and 34.69%, respectively. The model showed close agreement with the experimental treatment effects for pressure and for wall thickness, while stiffness was reduced more strongly in the simulations than in the experimental data. These results indicate that the calibrated framework can predict the direction and magnitude of therapy-induced improvements in MCT-induced PH while preserving the partial, rather than complete, reversal of disease remodeling.

### 3.5 Sensitivity of predicted treatment response to therapy parameters

We next examined the sensitivity of treatment predictions to the therapy-related parameters to better understand their time-dependent effects. Each parameter was varied independently while the others were held fixed. As shown in Fig. 8, all three therapy parameters influenced predicted remodeling outcomes, but with distinct patterns. Decreasing *ϕ_K_i__* reduced pressure, wall thickness, and stiffness, with 4-week pressure reduction ranging from 37.31% to 49.20%, thickness reduction from 10.35% to 36.86%, and stiffness reduction from 56.33% to 65.32%. In contrast, varying *ϕ_γ_* had a stronger effect on passive stiffening than on wall growth, with 4-week pressure reduction ranging from 14.53% to 53.10%, thickness reduction from 13.21% to 28.69%, and stiffness reduction from 12.63% to 77.08%. Thus, lowering *ϕ_γ_* produced the largest changes in stiffness and also substantially affected pressure. The activation timescale *τ*_Tx_ primarily controlled the rate at which treatment effects emerged. Smaller values of *τ*_Tx_ led to earlier divergence from the untreated trajectory and larger reductions by week 4. Across the tested range, pressure reduction varied from 26.47% to 42.41%, thickness reduction from 13.68% to 22.25%, and stiffness reduction from 33.87% to 60.08%. Compared with *ϕ_K_i__* and *ϕ_γ_*, variation in *τ*_Tx_ had a more modest effect on wall thickness but still meaningfully altered pressure and stiffness. Notably, we observed a distinct range of values that led only to an inflection in continued progression, where each of the metric values continued to increase or decreased for a short time before again increasing through the end of the simulation. This suggests that interventions must lead to functional changes of a certain threshold to interrupt or reverse disease progression.

**Fig. 8:**
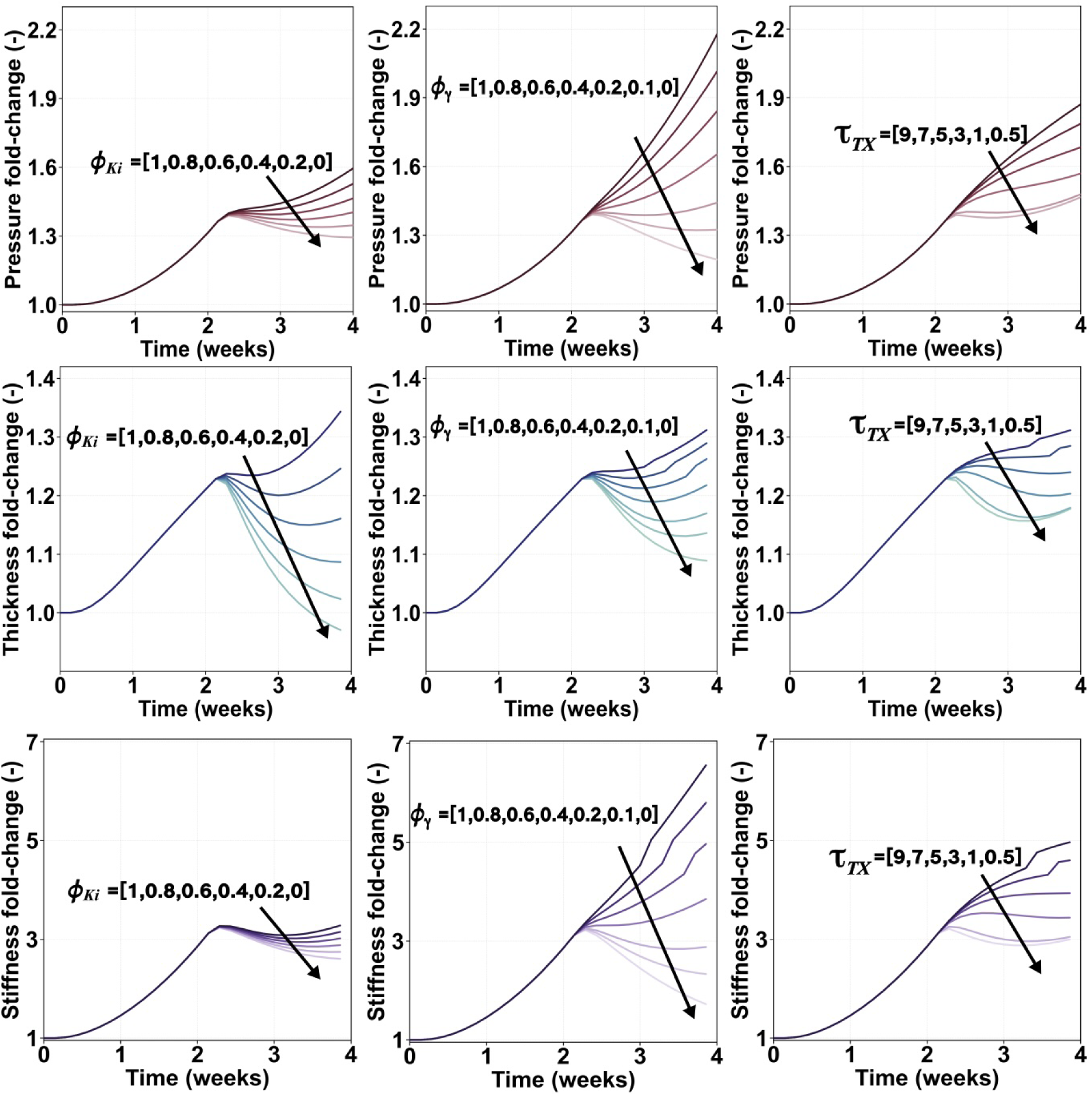
Sensitivity of predicted treatment response to therapy-related parameters. Foldchange trajectories of pressure (top), wall thickness (middle), and stiffness (bottom) are shown for independent variation of *ϕ_K_i__* (left), *ϕ_γ_* (center), and *τ* (right). Lower values of *ϕ_K_i__* and *ϕ_γ_* represent stronger suppression of maladaptive smooth muscle production and passive stiffening, respectively, while smaller *τ*_Tx_ values produce a faster treatment effect. The largest sensitivity to *ϕ_γ_* is observed in stiffness.

Overall, these results show that predicted treatment response is highly sensitive to how therapy is mapped onto maladaptive remodeling mechanisms. Reducing *ϕ_K_i__* primarily suppressed growth-related remodeling, reducing *ϕ_γ_* most strongly attenuated passive stiffening, and decreasing *τ*_Tx_ accelerated the onset of therapeutic benefit. These trends support the use of mechanistically defined therapy parameters for interpreting and predicting treatment response in MCT-induced PH.

## 4 Discussion

In this study, we developed a predictive pulmonary arterial G&R framework that couples morphometric tree hemodynamics with disease-specific maladaptive remodeling relations and therapy-modified intervention terms. Inflammatory parameters related to known pathophysiological changes in PH development were calibrated with numerical optimization to aggregated data from rat studies of PH induced by MCT and SuHx. Next, study-specific MCT and SuHx data on pressure, wall thickness, and stiffness were similarly captured. In doing so, we demonstrated the suitability of our constitutive assumptions to capture time course development of PH in a multiscale model focused on linking evolution of the vascular wall in the PA tree with organ-level hemodynamics.

To understand model sensitivity, we first performed parametric studies that clarified the distinct roles of the three maladaptive remodeling parameters. As shown in Fig. 3, increasing *K_i_* amplified overall remodeling severity across pressure, wall thickness, and stiffness, increasing *γ_i_* most strongly enhanced passive stiffening, and changing *δ_i_* primarily shifted the timing and rate of remodeling onset. This separation of effects is important because it shows that these few inflammatory parameters control much of the progressive evolution while remaining functionally interpretable in the context of aggregated cellular pathways, such as those commonly evaluated in gene ontology studies [9]. In particular, the dominant influence of *γ_i_* on stiffness is consistent with prior experimental and biomechanical studies suggesting that passive stiffening is an important biomarker for pulmonary vascular remodeling and pressure progression [4, 5, 18]. More broadly, these trends support the idea that PH progression is governed not only by geometric or topological changes, but also by evolution of wall composition that leads to positive feedback between biomechanical and inflammatory changes [22, 28].

Next, we examined our model’s ability to capture real time course data in PH progression. Aggregated pressure comparisons in Fig. 4 showed that the framework could reproduce the mean temporal progression of pressure across both MCT and SuHx datasets, despite heterogeneity across contributing studies and disease etiologies. In examining a richer dataset with multiple metrics comparable to model outputs, we reproduced the study-specific experimental MCT dataset reported by Liu et al. [18] with good overall fits across pressure, wall thickness, and stiffness. As shown in Fig. 5, the optimized model captured the progressive increase in all three outputs, with the strongest agreement observed for stiffness. The best fit parameter values further suggest that the MCT phenotype in this experimental dataset is characterized by strong maladaptive smooth muscle production, gradual remodeling activation, and passive stiffening. This demonstrates that the framework can move beyond qualitative markers of behavior and directly reproduce the measured trajectory of pulmonary vascular remodeling in experimental data.

At the same time, the study-specific SuHx results in Fig. 6 were much weaker than the corresponding MCT fit. MCT and SuHx represent different remodeling phenotypes and severities, with SuHx generally producing a more severe and partially irreversible form of PH with a time course that is distinct from the development of MCT-induced PH [17]. Therefore, the weaker SuHx fit suggests that the simple inflammatory constitutive formulation is sufficient to capture the trends in remodeling, but not the full mechanistic complexity of the more severe phenotype. In observing the time course from Liu et al [18]., we observed different trends in pressure and thickness vs stiffness, where pressure and thickness appear to plateau from 5 to 8 weeks, while stiffness continues to increase. As such, the SuHx results may point to the need for additional, more complex constitutive relations that can account for distinct time scales across processes like mass production and stiffening. Indeed, a key potential difference in MCT and SuHx is in the spatial effects of the initial insult, where MCT damages endothelial cells throughout the vasculature, and SuHx induces hypoxic vasoconstriction that varies across vessels of different caliber. Thus, a move towards spatially heterogeneous inflammatory stimuli may be necessary to capture the full breadth of PH phenotypes [15, 16].

With a calibrated model, we sought to identify its utility as a tool for simulating effects of therapy. As our goal was to show predictive capabilities, we chose to use in vitro data from PA SMC culture experiments to determine the magnitude of functional changes to inflammatory parameters. In Fig. 7, therapy reduced pressure, wall thickness, and stiffness relative to untreated MCT, with reasonable agreement for all metrics to measured outcomes for an equivalent experimental therapy [18]. As such, the current therapy formulation captures the direction of treatment benefit and the partial reversibility of remodeling, but may overestimate reversal of passive stiffness or fail to capture all mechanisms that maintain residual disease burden after treatment. The sensitivity analysis in Fig. 8 further clarified these mechanisms. Lowering *ϕ_K_i__* primarily suppressed growth-related remodeling, lowering *ϕ_γ_* produced the strongest effect on stiffness and also substantially affected pressure, and reducing *τ*_Tx_ accelerated the onset of treatment benefit. These results suggest that treatment response is not controlled by a single pathway and that hemodynamic improvement does not necessarily imply complete reversal of vessel remodeling. This interpretation is also consistent with the broader literature showing that advanced vascular remodeling can reduce reversibility and limit the effectiveness of vasodilatory therapies alone [5, 6, 11, 18]. Furthermore, these parametric studies demonstrated many cases without reversal of disease progression, as pressures continued to increase, albeit more slowly than in the case without therapy. Examining the ability of specific functional changes to fully reverse or interrupt disease progression is a critical use for this modeling approach that could guide the future development of protocols for clinical therapy administration.

Compared with the previous framework of Szafron et al. [6], the present study differs in several important ways. The prior work primarily established and explored a mechanistic framework for adaptive and maladaptive pulmonary arterial remodeling, whereas the present study directly calibrated the model to longitudinal experimental data and reported quantitative agreement for pressure, wall thickness, stiffness, therapy response, and aggregated cross-study pressure progression. While the earlier work established cumulative maladaptive constitutive changes, including smooth muscle hyperplasia, passive stiffening, synthetic phenotype, mechanobiological dysfunction, elastin degradation, and non-uniform WSS-mediated inflammation, the present study examined a subset of mechanisms in the disease formulation centered on maladaptive production, activation timing, and passive stiffening. Our earlier study focused strongly on establishing physiologic homeostatic behavior and demonstrating the necessity of hemodynamic feedback, whereas the present work shifts the main emphasis toward data-driven disease progression and intervention prediction. Although the previous study discussed potential treatment relevance and reversibility, they did not calibrate and test a specific therapy formulation against intervention data, whereas the present study implemented a treprostinil therapy formulation with explicit *ϕ_K_i__*, *ϕ_γ_*, and *τ*_Tx_ terms and evaluated predicted treatment response against reported therapy outcomes [18]. Thus, the present study includes extensive new work focused on quantitative methods for capturing disease progression and the constitutive modeling of therapy.

Several limitations should be acknowledged. First, remodeling was represented using order-averaged vessel behavior, which greatly improves computational tractability but reduces the ability to resolve spatial heterogeneity among vessels of the same order. Second, although the wall model contains multiple structurally important constituents, the disease-specific formulation focused mainly on smooth muscle-driven maladaptation and therefore does not yet fully represent the broader extracellular-matrix, endothelial, inflammatory, or elastin-related mechanisms that may be especially important in severe phenotypes such as SuHx. Third, the therapy analysis was based on relatively short-term intervention data, so long-term reversibility of the remodeled wall remains uncertain. Fourth, although LHS improved robustness of the inverse problem, practical identifiability remains a central challenge because the optimized parameters govern time-dependent biological responses rather than static baseline physiology [29]. Finally, the hemodynamic model remains a reduced-order representation with fixed morphometric structure and Poiseuille-type assumptions, which is appropriate for the present framework but does not capture all spatial and unsteady features of pulmonary vascular flow [6, 30].

For future work, the first priority is to refine the SuHx remodeling formulation by testing alternative constitutive relations, phenotype-specific stimulus functions, and additional optimized parameters in order to better represent more severe and partially irreversible disease, as well as enrich the model by incorporating varying inflammatory stimuli, pruning or structural rarefaction, right ventricular coupling, and higher-fidelity hemodynamics. A second priority is to extend the therapy framework to longer simulation horizons and to mechanism-isolation studies that independently suppress maladaptive production, passive stiffening, or both, thereby clarifying whether long-term recovery approaches baseline or instead plateaus above it. Another important direction is to expand the intervention framework to evaluate additional pulmonary hypertension therapies and drug effects beyond the current treatment case, allowing the model to compare mechanistically distinct therapeutic strategies within the same multiscale setting. Finally, improved uncertainty and identifiability analysis by quantifying parameter correlations, admissible solution ranges, and the measurements most needed to constrain the inverse problem would improve the utility in the model for decision making and driving future study design. More broadly, we will aim to preserve the mechanistic interpretability of the present framework while improving its ability to distinguish disease-specific pathways, predict reversibility, and support intervention design across different pulmonary hypertension phenotypes.

## 6 Supplement

### 6.1 Verification of the converted G&R framework under adaptive flow loading

We verified the converted Python framework by comparing its response to altered hemodynamic loading with the original MATLAB implementation reported by Szafron et al. [6]. An 80% increase in inlet flow was applied, and the resulting changes in normalized radius, normalized wall thickness, IMS, and WSS were compared across vessel orders. Fig 9A shows the results obtained using the original MATLAB implementation, whereas Figure 9B shows the corresponding results from the converted Python implementation. Vessel orders range from 1 to 10, with order 1 representing the smallest, most distal vessels and order 10 representing the largest, most proximal vessels. Increasing vessel order is indicated by the arrow in each panel. The Python implementation reproduced the order dependent temporal responses observed in the original MATLAB framework for all four quantities. In particular, the converted model captured the expected increases in vessel radius, changes in wall thickness, and transient adaptations in IMS and WSS following the increase in flow. This confirms that the coupled morphometric tree hemodynamics and G&R framework retained its adaptive behavior following conversion to Python and provides a verified foundation for introducing disease specific constitutive and inflammatory remodeling mechanisms.

**Fig. 9:**
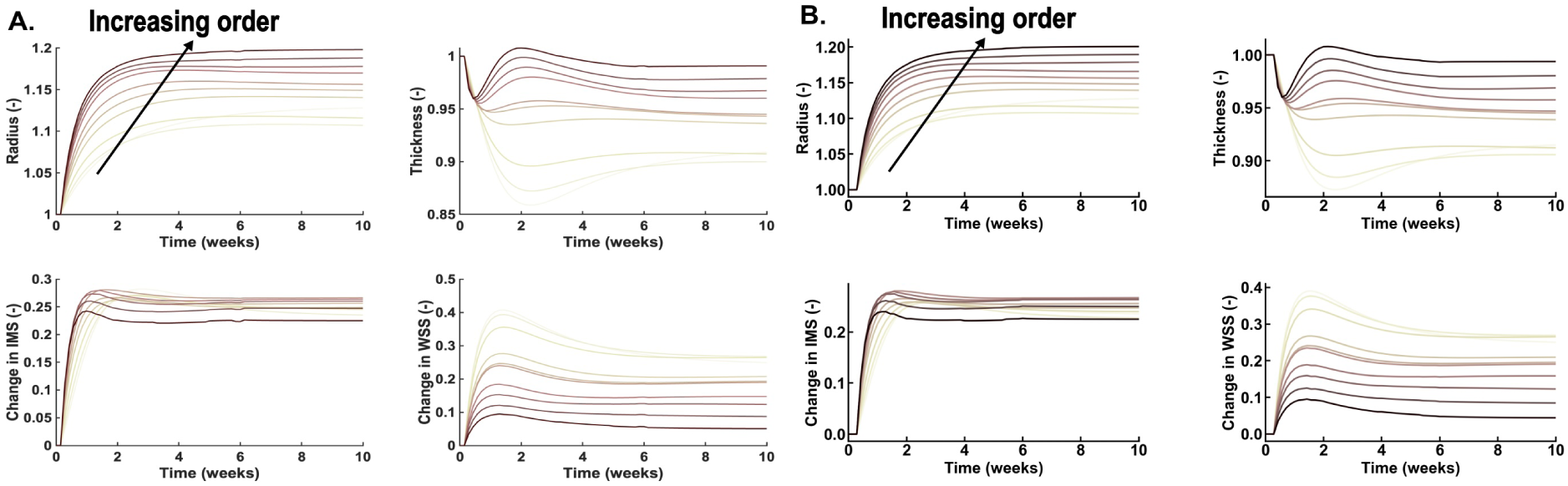
Verification of the converted G&R framework under adaptive flow loading. **(A)** Results from the original MATLAB implementation reported by Szafron et al. [6]. **(B)** Corresponding results obtained using the converted Python implementation. The plots show the order-dependent evolution of radius, wall thickness, IMS, and WSS following an 80% increase in inlet flow. Orders range from 1 (smallest, distal) to 10 (largest, proximal), with arrows indicating increasing order. This verification was performed in the default adaptive setting, prior to the introduction of any disease-specific inflammatory remodeling.

### 6.2 Data sources from literature on rat models of PH

**Table 2:** Studies contributing to the aggregated MCT pressure dataset.

| Paper | Time course (weeks) | Number of data points |
| --- | --- | --- |
| Liu et al., 2016 [18] | 0, 1, 2, 3, 4 | 5 |
| Zimmer et al., 2021 [31] | 0, 1, 2, 3 | 4 |
| Pankey et al., 2012 [32] | 0, 4 | 2 |
| Zaiman et al., 2008 [33] | 0, 4 | 2 |
| Mam et al., 2010 [34] | 0, 4 | 2 |
| Zhu et al., 2020 [35] | 0, 4 | 2 |
| Schwenke et al., 2009 [36] | 0, 3 | 2 |
| Chen et al., 2001 [37] | 0, 3 | 2 |
| Reindel et al., 1990 [38] | 0, 2 | 2 |
| Ruiter et al., 2013 [39] | 4 | 1 |
| Boueiz et al., 2009 [40] | 3 | 1 |
| Dorfmueller et al., 2011 [41] | 0 | 1 |
| Kosanovic et al., 2011 [42] | 0 | 1 |

**Table 3:** Studies contributing to the aggregated SuHx pressure dataset.

| Paper | Time course (weeks) | Number of data points |
| --- | --- | --- |
| Liu et al., 2016 [18] | 0, 1, 2, 5, 8 | 5 |
| Kato et al., 2017 [43] | 0, 1, 3, 5, 8 | 5 |
| Toba et al., 2014 [44] | 0, 1, 3, 5, 8 | 5 |
| Zungu-Edmondson et al., 2016 [45] | 0, 5, 8 | 3 |
| Chaudhary et al., 2018 [46] | 0, 4, 7 | 3 |
| Christou et al., 2018 [47] | 0, 7 | 2 |

### 6.3 Reduced-parameter optimization of aggregated pressure datasets

To evaluate the number of inflammatory remodeling parameters required to capture pressure progression, reduced-parameter optimizations were performed for the aggregated MCT and SuHx datasets. As shown in Supplemental Fig. 10A, optimization of *K_i_* alone accurately reproduced the mean MCT pressure trajectory over 4 weeks, yielding *R*^2^ = 0.9678 and RMSE = 0.0737. For the aggregated SuHx dataset, a one-parameter optimization using *K_i_* alone produced substantially weaker agreement with the experimental mean trajectory (*R*^2^ = 0.4795, RMSE = 0.6468; Supplemental Fig. 10B, left). Adding *δ_i_* as a second optimized parameter improved the SuHx fit to *R*^2^ = 0.8272 and RMSE = 0.3727 (Supplemental Fig. 10B, right), demonstrating that the temporal development of SuHx-induced pressure elevation could not be represented adequately through modulation of inflammatory magnitude alone.

**Fig. 10:**
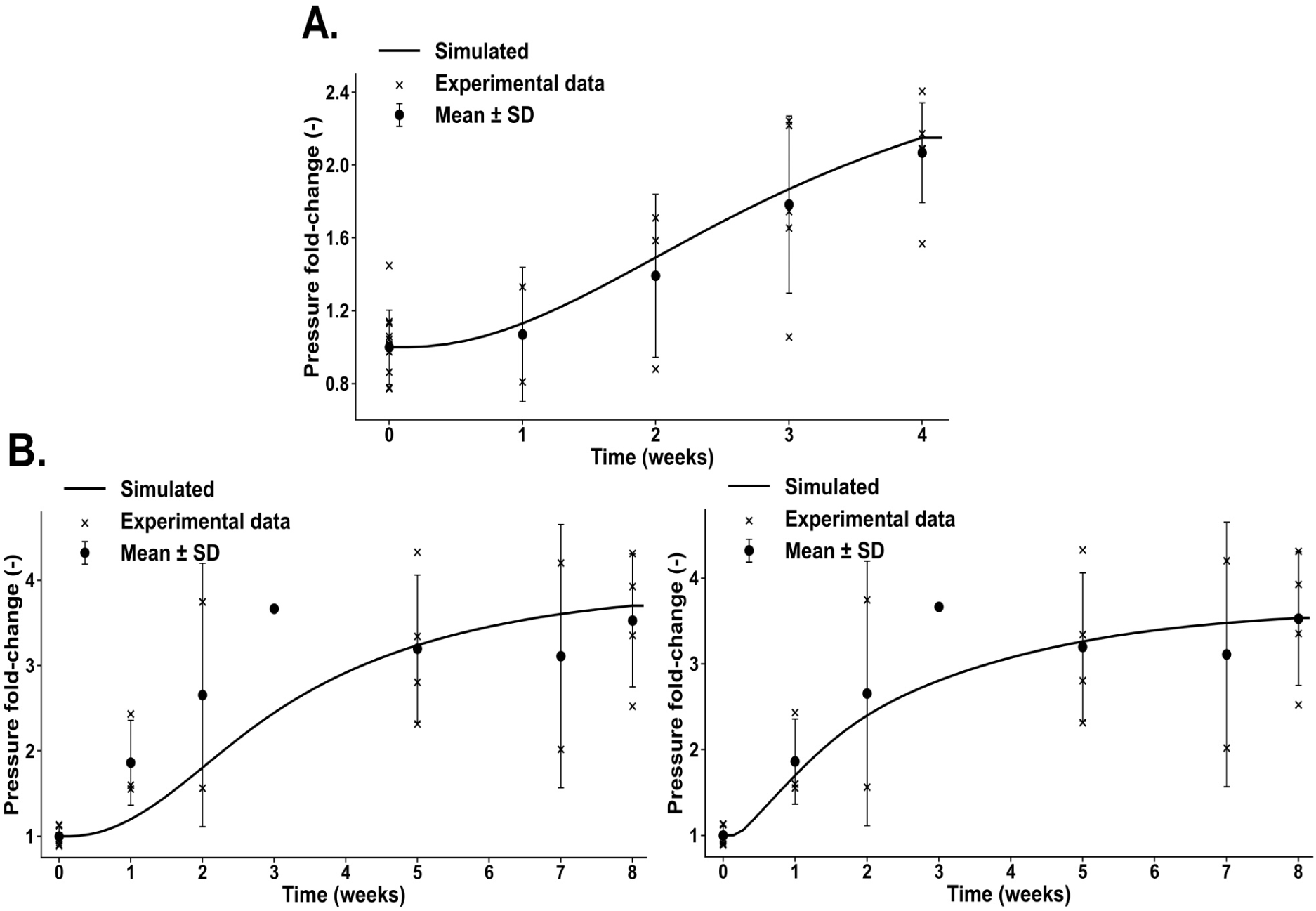
Reduced-parameter optimization of pressure progression across aggregated MCT and SuHx datasets. **(A)** Aggregated MCT pressure progression fitted by optimizing *K_i_* alone. The one-parameter model captured the mean experimental trajectory with *R*^2^ = 0.9678 and RMSE = 0.0737. **(B, left)** Aggregated SuHx pressure progression fitted by optimizing *K_i_* alone, yielding *R*^2^ = 0.4795 and RMSE = 0.6468. **(B, right)** Aggregated SuHx pressure progression fitted by optimizing *K_i_* and *δ_i_*, yielding *R*^2^ = 0.8272 and RMSE = 0.3727. Solid lines denote simulated pressure trajectories, crosses denote individual experimental measurements, and filled circles with error bars denote the experimental mean *±* SD at each time point. The improved SuHx fit obtained by including *δ_i_* demonstrates the importance of representing both the magnitude and temporal development of maladaptive remodeling.

### 6.4 Convergence of multi-objective optimization from multiple initial parameter sets

To evaluate the robustness of the study-specific multi-objective optimization, 15 initial parameter sets were generated using Latin hypercube sampling (LHS). Of these, nine optimization runs converged to similar parameter values and component cost-function values, whereas six converged to poorer local optima. Supplemental Fig. 11 shows representative optimization trajectories from two distinct initial parameter sets that converged to the same solution. Despite differences in their initial values and optimization paths, both runs approached the prescribed upper bounds for *K_i_* and *γ_i_* and a similar value of *δ_i_*, while producing nearly identical final pressure, thickness, and stiffness cost-function values.

**Fig. 11:**
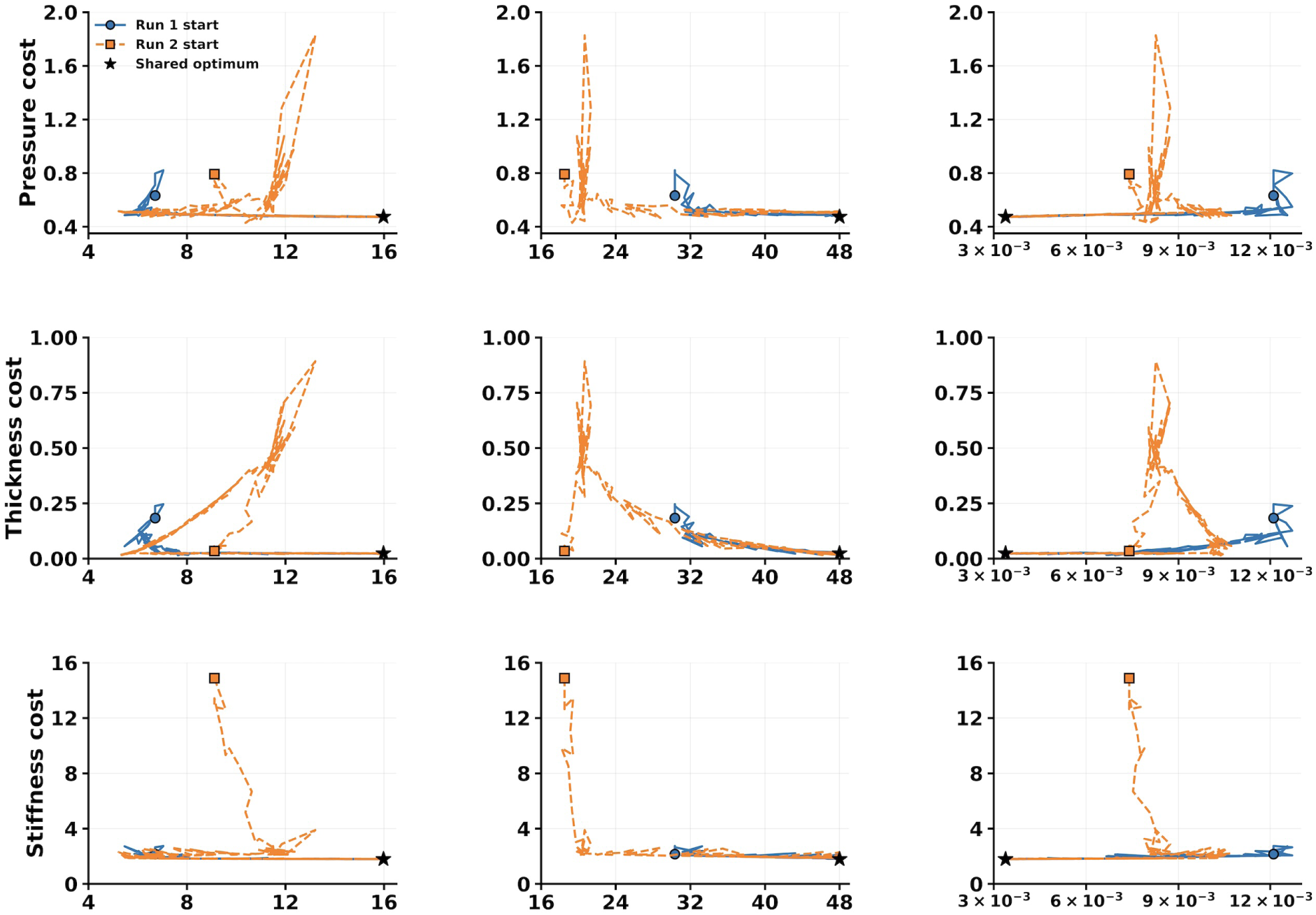
Convergence of repeated multi-objective optimizations from distinct initial parameter sets. Representative optimization trajectories for the pressure cost function (top row), thickness cost function (middle row), and stiffness cost function (bottom row) are shown as functions of *K_i_* (left column), *γ_i_* (middle column), and *δ_i_* (right column). The blue solid and orange dashed trajectories correspond to optimization runs initiated from two different parameter sets, with circles and squares indicating their respective starting points. The black star indicates the shared optimum approached by both runs. Although the trajectories followed different paths through parameter space, both runs converged to similar parameter and component cost-function values.

